## Supplemental Material 1 for "The effect of ethanol concentration on the morphological and molecular preservation of insects for biodiversity studies"

### Supplementary Material

Table Of Contents:


Quantification - spike-in plasmid 1
Amplicon data analysis pipeline 2
Table S1. 3
Table S2. 4
Table S3. 5
Table S4. 5


---

### Quantification - spike-in plasmid

The insert sequence was developed based on *Calliophora vomitoria* COI sequence, and synthesized and inserted into vector backbone pEX-A128 by Eurofins Genomics. The plasmid was cloned into JO-FI competent cells derived from the E. coli DH5α strain (A&A Biotechnology) using CloneJET PCR Clonning kit (Thermo Fisher Scientific). A single successfully transformed colony was transferred into LB medium (A&A Biotechnology) mixed with Ampicilin Sodium (A&A Biotechnology) at the final concentration 100 ug/ml and grown overnight at 37˚C. Plasmids were extracted from an overnight culture using Plasmid Miniprep DNA Purification Kit (EurX), linearized using single-cutting enzyme AatII (New England Biosciences), quantified using Qubit dsDNA HS Assay Kit (Thermo Fisher Scientific), and diluted with TE to the working concentration.

The insert sequence, with BF3/BR2 primer sites underlined:

>COI_CallioSynth

GGAGCCC<u>CAGATATAGCATTCCCTCG</u>AATAAAATTATTACATATATTTGCTATACTTAGTTCACTATCGTAA
GCATAAATATACAGAATATGAAATATTGGAGCTGGAACTGGATGAACTGTTTATCCACCTTTATCGCCATGC
ATAACTCTTCACTGATCTAAGGACGATATGGTTAGTATCTGTATACTTAGACGATGTATAAAGTCGAATCAA
TGCGTTAGAAACATTTATAGAATAGATAATTAGACATATTGTACGCCTAATAAGAACAAGATAAGACGCTA
AATCCAATCAATTCTTCAAGTAGTATACTATACTGTACTATATTCTTTACCAGTATTAGCAGGAGCTATTACT
ATATTTCGATTACCTCTAGTTATAATAAATGTAATGATAAACGAAGTAATACAAGTAGAACTATTCATATAA
CATGATTTATTT<u>TGATTTTTTGGTCATCCTGA</u>AGTTTAT

### Amplicon data analysis pipeline

The amplicon data were analyzed using Mothur v.1.44.2, using a custom pipeline. Commands are provided below.

```
   # Merging reads into contigs
make.contigs(file=DNA.txt)

   # variable-length insert primer trimming, screening contigs based on
   quality and length
trim.seqs(fasta=DNA.trim.contigs.fasta,
oligos=/mnt/matrix/symbio/db/references/primers_to_trim_VarLenIns.oligos,
processors=32, minlength=400, maxlength=430, maxambig=0, maxhomop=10,
pdiffs=2)
```

```
# primers_to_trim_VarLenIns.oligos ---
# oligos used for library prep:
# COIBF3_P5   ACACTCTTTCCCTACACGACGCTCTTCCGATCT---
   {insert}CCHGAYATRGCHTTYCCHCG
# COIBR2_P7   GTGACTGGAGTTCAGACGTGTGCTCTTCCGATCT---
   {insert)CDGGRTGNCCRAARAAYCA

forward CCHGAYATRGCHTTYCCHCG
forward ACCHGAYATRGCHTTYCCHCG
forward GACCHGAYATRGCHTTYCCHCG
forward TGACCHGAYATRGCHTTYCCHCG
reverse CDGGRTGNCCRAARAAYCA
reverse TCDGGRTGNCCRAARAAYCA
reverse ATCDGGRTGNCCRAARAAYCA
reverse GATCDGGRTGNCCRAARAAYCA
```

```
   # selecting only the sequences that survived the screen from the group
file
list.seqs(fasta=DNA.trim.contigs.trim.fasta)
get.seqs(accnos=DNA.trim.contigs.trim.accnos, group=DNA.contigs.groups)

   # read dereplication
unique.seqs(fasta=DNA.trim.contigs.trim.fasta)
count.seqs(name=DNA.trim.contigs.trim.names,
group=DNA.contigs.pick.groups, compress=f)

   # discarding singletons
split.abund(fasta=DNA.trim.contigs.trim.unique.fasta,
count=DNA.trim.contigs.trim.count_table, cutoff=1)

   # aligning reads against a custom reference - alignment of previously
   obtained COI sequences of the experimental species, the spike-in
   standard, and alphaproteobacterial symbionts
align.seqs(fasta=DNA.trim.contigs.trim.unique.abund.fasta,
reference=/mnt/matrix/symbio/db/references/COI_ref_seqs_fragility.fasta,
processors=40)
```

```
   # discarding unaligned or poorly aligned sequences, filtering the
   alignment
screen.seqs(fasta=DNA.trim.contigs.trim.unique.abund.align,
count=DNA.trim.contigs.trim.abund.count_table,            minlength=410,
maxlength=425, start=2, end=430)
filter.seqs(fasta=DNA.trim.contigs.trim.unique.abund.good.align,vertical=T
, trump=.)

   # clustering, using the Vsearch agc algorithm
cluster(fasta=DNA_final.fasta,   count=DNA_final.count_table,   method=agc,
cutoff=0.03)

   # based on clustering results, binning sequences, generating OTU table,
   and identifying representative sequences for OTUs
bin.seqs(list=DNA_final.agc.list, fasta=DNA_final.fasta, label=0.03)
make.shared(list=DNA_final.agc.list,          count=DNA_final.count_table,
label=0.03)
get.oturep(list=DNA_final.agc.list,        fasta=DNA_final.agc.0.03.fasta,
count=DNA_final.count_table, method=abundance, cutoff=0.03)
```

**Table S1.**

Species included in the communities, number of individuals used, expected brittleness, and origin.

| Species | No. individuals | Brittleness | Origin |
|---|---|---|---|
| *Macrolophus pygmaeus* | 10 | Weak | Commercial purchase: Lindesro AB (Sweden) |
| *Aphidoletes aphidimyza* | 10 | Very weak | Commercial purchase: Lindesro AB (Sweden) |
| *Drosophila hydei* | 10 | Tough | Commercial purchase: Fibe AB (Sweden) |
| *Dacnusa sibirica* | 10 | Tough | Commercial purchase: Lindesro AB (Sweden) |
| *Calliphora vomitoria* | 2 | Intermediate | Commercial purchase: Reptilgrottan (Sweden) |
| *Formica rufa* | 2 | Very tough | Collected from NRM surroundings |
| *Dermestes haemorrhoidalis* | 2 | Very tough | Donated from NRM vertebrate collection |

**Table S2.**

Pairwise comparisons for treatments with significant differences in Experiment 1. Treatment 1 is different from Treatment 2, being the mean number of appendages lost higher/lower in Treatment 2. Number = ethanol concentration, G = Gentle shaking, V= Violent shaking.

| Species | Treatment 1 | Treatment 2 | Lower / Higher |
|---|---|---|---|
| *Macrolophus pygameus* | 30G | 80G, 90G, 95G, 97G | Higher |
| | 30G | 70V, 90V, 95V, 97V, 99V | Higher |
| | 50G | 95V | Higher |
| | 95G | 30V | Lower |
| | 30V | 95V | Higher |
| | 50V | 95V | Higher |
| | 80V | 95V | Higher |
| *Aphidoletes aphidimyza* | 30G | 90G, 95G, 97G, 99G | Higher |
| | 30G | 90V, 95V, 97V, 99V | Higher |
| | 50G | 90G, 95G, 97G, 99G | Higher |
| | 50G | 90V, 95V, 97V, 99V | Higher |
| | 50G | 30V | Higher |
| | 70G | 90G, 95G, 97G, 99G | Higher |
| | 70G | 30V | Higher |
| | 70G | 90V, 95V, 97V, 99V | Higher |
| | 80G | 90G, 95G, 97G, 99G | Higher |
| | 80G | 95V, 95V, 99V | Higher |
| | 90G | 30V, 50V, 70V, 80V | Lower |
| | 90G | 99V | Higher |
| | 95G | 30V, 50V, 70V, 80V | Lower |
| | 97G | 30V, 50V, 70V, 80V, 90V | Lower |
| | 99G | 30V, 50V, 70V, 80V, 90V | Lower |
| | 30V | 95V, 97V, 99V | Higher |
| | 50V | 95V, 97V, 99V | Higher |
| | 70V | 90V, 95V, 97V, 99V | Higher |
| | 80V | 95V, 97V, 99V | Higher |
| | 90V | 97V, 99V | Higher |
| *Drosophila hydei* | 50G | 90V | Lower |
| | 97G | 90V | Lower |
| | 30V | 90V | Lower |
| | 50V | 90V | Lower |
| | 70V | 90V | Lower |
| *Calliphora vomitoria* | 70G | 95G, 99G | Higher |
| | 70G | 95V, 99V | Higher |
| | 70V | 95G, 99G | Higher |
| | 70V | 95V, 99V | Higher |

**Table S3.**

Pairwise comparisons for treatments with significant differences in Experiment 2. Treatment 1 is different from Treatment 2, being the mean number of appendages lost higher/lower in Treatment 2. Number = ethanol concentration, W = Walking transport, R= Running transport, PN = PostNord transport.

| Species | Treatment 1 | Treatment 2 | Lower / Higher |
|---|---|---|---|
| *Macrolophus pygameus* | 70W | 70R, 95R | Higher |
| | 70W | 70PN, 95PN | Higher |
| | 95W | 70R, 95R | Higher |
| | 95W | 70R, 95R | Higher |
| | 70R | 95R | Higher |
| | 95R | 70PN | Lower |
| | 70PN | 95PN | Higher |
| *Aphidoletes aphidimyza* | 70W | 95R | Higher |
| | 70W | 95PN | Higher |
| | 95W | 95R | Higher |
| | 95W | 95PN | Higher |
| | 70R | 95R | Higher |
| | 70R | 95PN | Higher |
| | 95R | 70PN | Lower |
| | 95R | 95PN | Lower |
| | 70PN | 95PN | Higher |

**Table S4.**

Pairwise comparisons for treatments with significant differences in Experiment 3. Treatment 1 is different from Treatment 2, being the mean number of appendages lost higher/lower in Treatment 2. Number = ethanol concentration, C = Control, D= Drying pre-treatment, F = Freezing pre-treatment.

| Species | Treatment 1 | Treatment 2 | Lower / Higher |
|---|---|---|---|
| *Macrolophus pygameus* | 70D | 70C, 95C | Higher |
| | 70D | 95D | Higher |
| | 95D | 70F, 95F | Higher |
| *Aphidoletes aphidimyza* | 70C | 95C | Higher |
| | 70C | 95D | Higher |
| | 70C | 95F | Higher |
| | 70D | 95C | Higher |
| | 70D | 95D | Higher |
| | 70D | 95F | Higher |
| | 70F | 95C | Higher |
| | 70F | 95D | Higher |
| | 70F | 95F | Higher |
| *Drosophila hydei* | 95D | 70F | Lower |
| *Calliphora vomitoria* | 70F | 95F | Higher |
